# Assessing ‘larger-than’ rule learning in African grey parrots (*Psittacus erithacus*)

**DOI:** 10.64898/2026.10.19.760036

**Authors:** Pizza Ka Yee Chow, Awani Bapat, Theo Robert, James R Davies, Auguste M P von Bayern

## Abstract

Many animals discriminate between magnitudes (e.g., in size, quantity, volume, length) that have fitness implications. Accumulated experience may facilitate the formation of relational rules that can be applied in novel contexts. However, the extent to which animals can form a relational rule, and are able to apply it to novel contexts remains a core focus when understanding the evolution of cognition. Here, we assessed African grey parrots’ ability to acquire the relative ‘larger-than’ rule in *size* or *quantity* discrimination, and if so, how this was learned. Eight captive parrots initially participated in a natural preference test (NPT) to assess their prior preference for a larger amount of seeds. Each parrot was then randomly selected to either acquire the larger-than rule in size or quantity in a basic training (BT). They then participated in transfer training (TT) to evaluate their ability to apply the learned rule to novel stimulus pairs, and a final transfer test (FTT) to assess whether the parrots could generalise the learned rule to unseen stimulus pairs beyond the training range. Six parrots successfully completed both training and the test, with only one parrot, Sensei, passing the quantity discrimination FTT. More detailed analyses showed that the parrots showed varied performance in the TT, but as a group, showed limited transfer in the TT, suggesting the use of associative learning through increased experience in BT. They also showed better performance if a stimulus pair had a larger difference (Weber’s law). Sensei’s success was attributed to extensive TT alongside his quick natural preference toward larger amounts. These results suggest that repeated exposure to TT stimulus pairs increases familiarity with the stimulus pairs, which can facilitate task success, but individuals vary in their ability to acquire relational rules.

## Introduction

Understanding relational concepts, such as ‘larger than’ or ‘smaller than’, involves mentally representing an abstract relationship between two objects along a shared dimension (e.g., size, quantity, volume, length). A prerequisite for such concepts requires individuals to be able to discriminate between varying magnitudes, or form a relational rule between objects, which many animals can do (Worsley et al. 2025a) e.g. primates (Anderson et al. 2005), birds (Garland et al. 2012; Brown et al. 2022), fish (Agrillo and Bisazza 2017; Miletto Petrazzini and Brennan 2020), insects (Pahl et al. 2013; Avarguès-Weber et al. 2014), reptiles (Agrillo 2014; Miletto Petrazzini et al. 2017), ungulates (Schaffer et al. 2025). Subsequently, animals may form relational concepts, whereby they apply the learned relational rule to novel contexts or across other dimensions. Being able to discriminate between varying magnitudes may allow individuals to flexibly optimise decision-making in different contexts, such as foraging, navigation, predator avoidance, and collective decision-making; see review by (Nieder 2020a; Worsley et al. 2025a). Such discrimination does not necessarily lead to relational rule learning or relational generalisation. However, being able to generalise a learned rule to novel contexts has been shown to have a neural basis that is favoured by evolution for species with high probabilities of encountering novel contexts, or to minimise costs of trial-and-error learning (Ghirlanda and Enquist 2003; Zentall et al. 2008). Given that relational concept learning is seen as a cornerstone of human cognition, understanding to what extent animals can demonstrate such concepts remains a central investigation in the evolution of cognition.

A typical experimental paradigm that is used to assess the ability to learn relational rules and to form relational concepts starts with presenting a pair of stimuli to individuals varying only in magnitude on the same dimension (e.g., size, quantity, volume). One of the stimuli is rewarded, and the other is not. Individuals acquire this reward contingency until they reach a stringent criterion reliably (e.g., ≥ 80% correct responses across two consecutive sessions).

Then the experiment proceeds to a transfer test. To assess relational rule learning, a transfer test includes a stimulus pair with one familiar (learned/old) stimulus and a novel stimulus, and again, both stimuli vary in magnitude on the same dimension (e.g., Lazareva 2012).

Evidence of relational rule learning can be demonstrated when individuals consistently apply the learned relational rule (e.g., ‘choose the larger’) to novel stimuli rather than merely responding to previous reinforcement. For example, the rule is learnt when the animal is consistently choosing the larger stimulus even when the previously rewarded stimulus is no longer the larger of the two. To assess relational concept formation, a transfer test includes completely new stimuli that can go beyond the dimensions of the stimuli used in training (e.g., size→length). Evidence of relational concepts exists when individuals apply the learned rule to other dimensions (i.e., choose whichever is larger when encountering novel stimuli of a different dimension, e.g. Miletto Petrazzini and Brennan 2020).

Individuals may vary in understanding hidden relational rules about stimulus relationships, and in generalising the rules to novel contexts. The acquisition of relational rules and concepts develops gradually; for example, human children demonstrate the ability to discriminate ‘more’ as early as three years old (Odic et al. 2013), and build on this to grasp the concept of ‘more than/larger than’, but may not fully understand the ‘larger-than-large’ concept even after becoming adults (Falk 1994; Hannula et al. 2006). In relational rule learning tasks, an increased experience with the training set may facilitate the formation of a rule and the generalisation of relational rules to novel contexts (Zentall et al. 2014a).

However, individuals can also use other strategies (e.g., associative learning) to discriminate between magnitudes during training. Thus, the success of generalisation may not only depend on whether a relational rule has been acquired, but also on prior training history and the similarities between the training and testing stimuli (Lazareva 2012). Performance in relational rule learning tasks across many animal species may also be affected by the perceptual discriminability of the differences between stimuli, also known as Weber’s law (Worsley et al. 2025b).

Our goal was to investigate to what extent African grey parrots are able to acquire the ‘larger-than’ rule and to extrapolate this rule to novel contexts. Two groups of parrots were trained to discriminate one of the two different dimensions (size or quantity) varying in magnitude.

Note that our interest was neither in parrots’ numerical sense nor in their quantification system, which has been shown (e.g. Pepperberg 1987a, 1994, 2006a). Instead, we assessed their performance based on their understanding of relations between the stimuli. African grey parrots are among the many bird species that have demonstrated advanced learning abilities across different cognitive tasks, such as match-to-sample (Pepperberg 1987b, 2006b; Pepperberg and Gordon 2005; Mikolasch et al. 2011; Pepperberg et al. 2013), inferential understanding of quantity and volumes (Pepperberg et al. 2017; Cornero et al. 2020), and demonstrated relative class concepts in size and quantity (Pepperberg and Brezinsky 1991a; Pepperberg 2006c; Aïn et al. 2009a). Despite this, it remains unclear whether African grey parrots can generalise such relational learning as an abstract “larger-than” rule across different contexts.

We first used a natural preference test (NPT) to assess whether the parrots have a prior preference for larger amounts of natural food, as well as to prompt them to choose the ‘larger-than’ food patch. The parrots were then randomly assigned to one of the two basic training (BT, see below for details) sessions, during which the larger of the two stimuli in the pair (size or quantity) was rewarded. Then they proceeded to the transfer training (TT), during which they were confronted with a set of three types of stimulus pairs: old-old stimulus pairs had both stimuli presented within the range of BT, but in novel shapes; old-new had one of the two stimuli presented beyond the range of BT in novel shapes, and new-new stimulus pairs had both stimuli completely beyond the range of BT and in novel shapes. Finally, the parrots experienced a final transfer test (FTT) in which the stimulus pairs included only novel stimuli beyond training ranges. If the parrots had formulated the relational concept, then they would be able to solve the old-new and new-new stimulus pairs in TT as well as the completely novel stimulus pairs in FTT beyond chance levels.

## Methods

### Parrots and housing

Eight captive African grey parrots (*Psittacus erithacus,* six females and two males), housed in a single semi-natural aviary at the Loro Parque Animal Embassy Zoo, participated in this project. The parrots were fed fruit twice a day, once in the morning and once in the evening. Seeds were also provided to them depending on their individual weights and accounting for nut rewards provided during the experiment. Water was provided *ad libitum.* For more details on housing for the parrots, see (Krasheninnikova et al. 2018). We tested the parrots 2 hours after the morning feeding, and no later than an hour before the evening feeding to maximise motivation. All individuals had training experience of being transported to the test room using a small portable cage (36 cm length x 31 cm width x 31 cm height); individual parrots walked into the cage voluntarily.

### Experiment set-up

The experiment involved a two-room set-up; the two rooms were connected but separated by a transparent glass window (89.5 x 88 cm, length x width). One room was for the parrot, and the other room was for the experimenter (Figure 1a). In the parrot’s room, there was a table (154.5 x 55 cm) positioned just next to the glass. A perch (15 x 5 x 6.5 cm, length x width x thickness) was fixed at the centre of the far edge of the table, approximately 55 cm away from the window; this allowed the parrots to see the seeds/stimuli from the perch (equidistant to the choices during the experiment). In the middle of the lower part of the window, there was a portion cut out and covered by a detachable transparent plexiglass (60 x 25.3 cm, length x width). The plexiglass had two holes (5 cm diameter, 23.1 cm apart) that were positioned at the eye level of the parrots so that they could put their head through the plexiglass during the experiment to indicate their choice. Between the plexiglass and at the bottom of the window, there was a small gap for reward delivery (only for the natural preference test). In the experimenter’s room, there was a wooden board (60 x 39.5 cm, length x width) positioned at the middle-bottom next to the window. Two A4 (29.7 x 21 cm) white plexiglass sheets were placed on the board in front of the two holes of the detachable plexiglass. All seeds/stimuli were presented on the white plexiglass sheets (to maximise saliency), corresponding to the front of the two holes. An opaque cardboard (98.5 cm x 44 cm) was used to cover the plexiglass window during the inter-trial intervals, so that the rebaiting process was not visible to the parrots.

**Figure 1.**
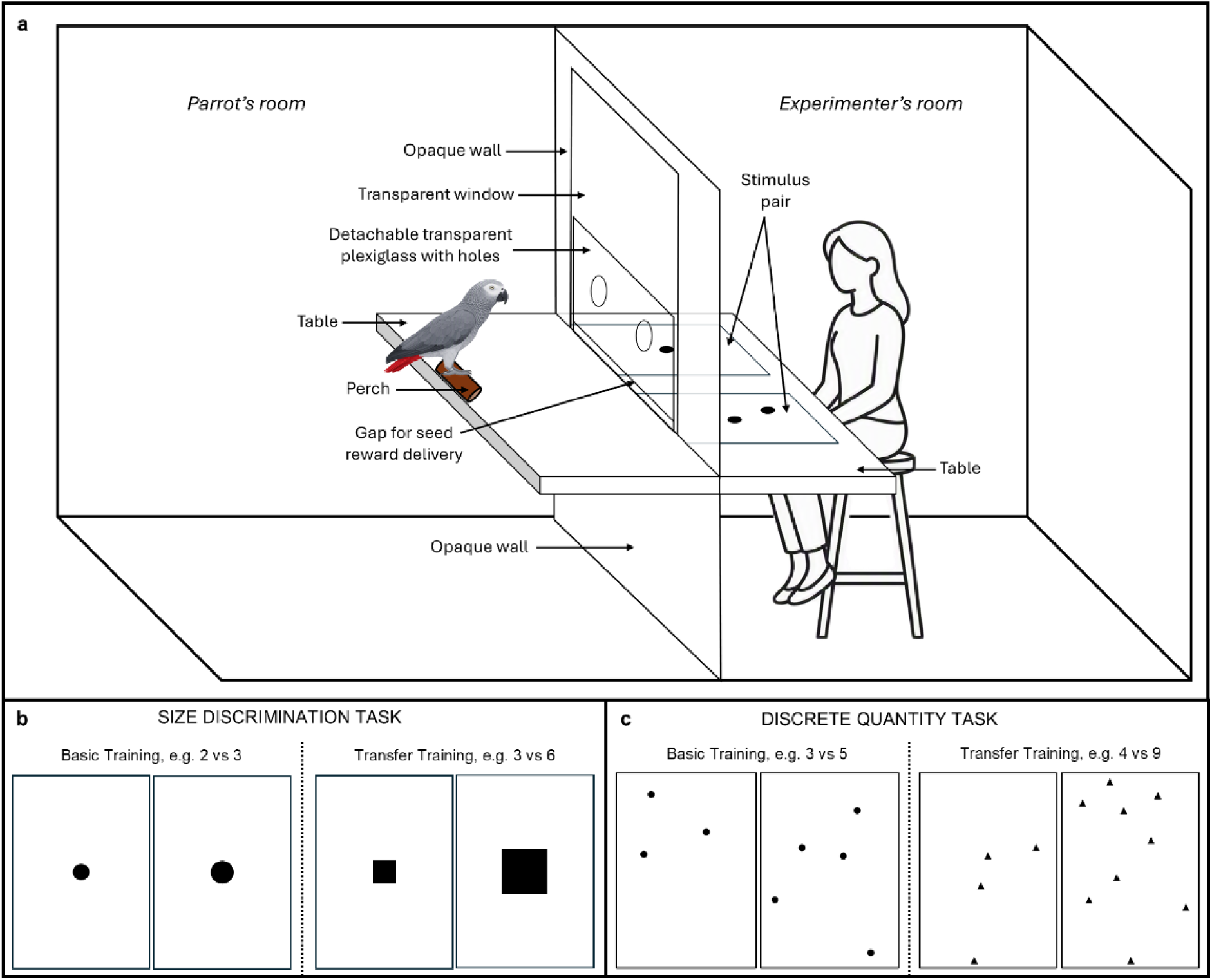
(a) Layout of the experiment compartments. (b) Examples of stimulus pairs used for the basic training (circles) and transfer training (squares) in the size experiment. (c) Examples of stimulus pairs used for the basic training (circles) and transfer training (triangles) in the discrete quantity experiment.

#### General procedures

A previous study with the same individual parrots used selective pairing with quantities ranging from 0 to 8, using sunflower seeds as rewards (Krasheninnikova et al. 2019). However, the pairings were selective, and thus we first conducted a spontaneous discrimination task (hereafter, natural preference test) for three purposes: first, to systematically and comprehensively assess whether parrots show a natural preference for larger quantities; second, to assess whether their preference was influenced by the previous study; and finally, to prompt the parrots to choose larger quantities. To achieve these purposes, we included all possible combinations of pairings to systematically assess parrots’ natural preference toward larger quantities. The natural preference test (NPT) consisted of 15 trials in each session, which included 10 trials with various pairs of unequal amounts of seeds (that is, different stimulus pairs, e.g., 1 vs 5, 2 vs 4), and 5 trials with pairs of equal amounts of seeds (that is, the same stimulus pairs, e.g., 1 vs 1, 2 vs 2). The same stimulus pairs were used to examine parrots’ side preference/bias. In each trial, a pair of seed combinations ranging from 1-5 were placed at the centre of each sheet. A parrot had to indicate his/her choice by inserting his/her head into one of the two holes corresponding to the sheet. Once the parrot indicated his/her choice, we slid the plate through the hole and the parrot could consume the corresponding chosen amount of seeds. After which, the experimenter used the opaque barrier to cover the hole while the parrot returned to the perch for the next trial (inter-trial interval 1 minute); this aimed to ensure the next trial started with presenting both stimuli equidistant to the parrot. The parrot completed one session (less than 30 minutes) per day until s/he showed a preference for a larger quantity (8 out of 10 different-size stimulus pairs in three out of five sessions) (see Results for more details).

Once all parrots had completed the NPT, we randomly assigned each parrot to either the size discrimination task (hereafter, ‘SDT’, Figure 1a) or the discrete quantity discrimination task (hereafter, ‘QDT’, Figure 1b). The procedure of the discrimination task was the same as the NPT. Each discrimination task consisted of three phases: basic training (BT), transfer training (TT), and a final transfer test (FTT). In all training tasks and the test, we first gave the parrots three ‘warm-up’ trials that included a medium-sized sunflower seed placed in the middle of each plate. No stimulus pairs were exposed to the parrots in these warm-up trials. The parrots had to consume both sunflower seeds before the next trial began; this ensured that they understood that both holes were available and that they familiarised themselves with the testing procedure.

For both tasks, the BT included 15 trials per session, with 10 different-size/number stimulus pairs and five same-size/number stimulus pairs. The same-sized stimulus pairs were presented in a mix with the different-sized stimulus pairs; they were used to examine parrots’ side preference/bias and served as an inter-trial reinforcement to keep the parrots’ motivation. The parrot had to indicate a choice by inserting its head through either hole, same as during the NPT. For the same stimulus pairs, they received a reward (a piece of walnut) for choosing either side. For the different stimulus pairs, they received a reward if they chose the larger-size stimulus, whereas choosing the smaller-size stimulus was unrewarded. In each session, we pseudo-randomised the presentation of stimulus pairs; the larger-size stimuli were shown equally on the left side and the right side. We ensured that the ‘larger-than’ stimuli did not appear on either the left or right side more than three consecutive times. Once the parrots reached the training criteria (i.e. 8 or more correct choices out of 10 for three consecutive sessions and no side bias shown), they participated in the TT.

In the TT, each training session consisted of 20 trials without same-size/number stimuli. Training stimulus pairs were novel to parrots and consisted of three types: (a) old-old pair, where novelty was in shapes and not numbers (e.g. 1 vs 3), (b) old-new pair, where novelty was in both shapes and one of the numbers was previously shown and the other was not shown in the training phase (e.g. 4 vs. 7), and (c) new-new pair, where novelty was in both shape and both numbers that were not shown in the training phase (e.g. 6 vs.10). Each of these pairs allowed us to analyse whether the parrots would be able to transfer the rule to novel stimuli based on the shape from the old-old stimulus pairs, to use a ‘more-than’ concept with a novel shape and an unseen numbers, and to use a ‘larger-than-large’ concept when a novel shape and two unseen numbers were presented to them. Similar to BT, the ‘larger-than’ stimuli (correct choice) were reinforced, whereas ‘smaller-than’ stimuli (incorrect choice) were not reinforced. We pseudo-randomised the sequence of presentation pairs so that the ‘larger-than’ stimuli were equally shown on the left and on the right side. The criterion for TT was 18 out of 20 trials (90%) in two consecutive sessions. Once a parrot passed the criterion, it went to the final transfer test that had 20 trials with unfamiliar stimulus pairs ranging from 11-20 cm (Figure 1c), either in absolute or total area.

In all training and tests, a session lasted no longer than 30 minutes. We minimised cueing individuals’ responses by wearing a pair of reflective sunglasses. Hand movements, however, were necessary, including removing and placing the barrier when a trial started and ended, respectively. Despite this, the hand that was used to move the barrier was randomised. We also pulled back stimuli after the individuals made a choice using the same hand as the one used to deliver rewards.

### Stimulus pairs

In the SDT, the stimulus was positioned at the centre of the A4 white plexiglass sheets (see Figure 1b). A dark circle (ranging from 1-5 cm diameter) was used for BT, whereas a dark square was used for TT (side ranging from 1-10 cm) and FTT (side ranging from 11-20 cm). In the QDT, the BT stimuli were dark circles (1 cm diameter) varying in number from 1 to 5, placed randomly on an A4 white paper (Figure 1c), with the placement never being repeated within or across sessions. Dark triangles (base and height 1cm) were used as the stimuli for TT (varying in number from 1 to 10) and FTT (varying in number from 11 to 20). For both experiments, a total of 25 pairs, including same-number/size stimuli (e.g. 1 vs. 1 cm, 2 x 2 cm) were generated for the BT, and 90 stimulus pairs (excluding same-size stimuli) were generated for the TT and FTT. In addition, the difference between two numbers () was calculated in each session for subsequent analyses.

### Data analysis

All eight parrots participated in the NPT. Three parrots (Jack, Jelo and Nikki) participated in the BT, TT and FTT of the SDT. Five parrots participated in the QDT. Of these, however, only three parrots (Kimmi, Lizzy and Sensei) completed BT, TT and FTT. The remaining two parrots were excluded from further training: Nina, who developed extreme side bias (87% in 70 sessions) and Bella, who was suspected of having health issues after 93 sessions. Thus, all subsequent statistical analyses were conducted only for the six birds who completed all stages of the study. All videos were recorded using two cameras, one in the parrot’s room and one in the experimenter’s room. The parrots’ choices were directly noted during the experiment. 10% of all sessions were coded by a second coder (J.R.D.) to assess inter-observer reliability.

All statistical analyses were conducted in R (version 4.5.2; R Core Team, 2025) using RStudio (version 2025.09.2, Posit team, 2025). Inter-observer reliability was assessed using the function ‘kappa2’ of the package ‘irr’ (version 0.84.1, Gamer et al. 2019), and the average Cohen’s Kappa was 0.94 for the three pairs of coders. To assess learning and performance of the parrots, four generalised linear mixed models (GLMMs) and two generalised linear models (GLMs) were fitted using the function ‘glmmTMB’ of package ‘glmmTMB’ (version 1.1.14, Brooks et al. 2017).

For each stage of the experiment, i.e. NPT (model 1), BT (model 3), TT (model 5) and FTT (model 6), the choice made by the parrot in each trial (1 = larger quantity, 0 = smaller quantity) was modelled using a GLMM with a binomial family distribution and logit link (Mccullagh and Nelder 1989). To examine whether the parrots learned to choose the larger quantity across the NPT, BT and TT sessions, Models 1, 3 and 5 included the session number as a fixed predictor. This fixed effect was not included in Model 6, as the FTT comprised only one session. To examine whether the parrot’s choice is influenced by Weber’s law, i.e., the perceptual contrast between stimuli, we included the ratio between the number or size metric of the stimuli (hereafter “stimulus pair ratio”) as a test fixed predictor in all four models. For the TT, where some of the stimulus pairs were novel to the parrots, Model 5 also included a fixed test predictor for stimulus pair type with levels “old-old”, “old-new”, and “new-new”. In the same model, we included the number of BT sessions required by each parrot as a test fixed predictor to see whether the experience in BT primes them to choose the larger quantity in TT. Similarly, model 6 included the number of NPT sessions, number of BT sessions, and number of TT sessions as fixed predictors. In Models 3, 5 and 6, we controlled for the fixed effect of experiment type (SDT vs QDT). To avoid pseudo-replication, all four models also included the random effect of individual identity (Schielzeth and Forstmeier 2009; Barr et al. 2013). In addition, Models 1, 3 and 5 also included the random effect of session number.

To assess whether the number of NPT sessions experienced affected the number of BT sessions required by each parrot, we fitted a GLM with a negative binomial family and logit link function (Model 2). Similarly, we fitted another negative binomial GLM to see whether the number of NPT and BT sessions experienced predicted the number of TT sessions required by each parrot (Model 4). In both models, experiment type was included as a control fixed predictor.

For model diagnostics of all models, we assessed the collinearity between fixed effect using the function ‘vif’ of package ‘car’ (version 3.0-6) (Fox and Weisberg 2018) and detected no issues (max VIF, model 1: 1.0, model 2: 1.4, model 3: 1.0, model 4: 2.3, model 5: 1.3, model 6: 3.7). For the GLMMs, we also examined the distribution of the best linear unbiased predictors (BLUPs) and found that all BLUPs were normally distributed with low variation within the levels of the random effects. For the GLM, we assessed the overdispersion of the model fit using the function ‘testDispersion’ of package ‘DHARMa’ (version 0.4.7, Hartig 2024) and found that the models were not overdispersed (dispersion parameters, model 2: 1.11, model 4: 1.16). The model stability of GLMs was assessed by computing the dfbeta values (Quinn and Keough 2002), whereas for the GLMMs we used the function ‘influence.me’ of the package ‘glmmTMB’.

Model inference was conducted by comparing the full models with the corresponding null models (including only the control fixed effects and random effects) using a likelihood ratio test (LRT, Dobson 2001) with the function ‘anova’. We tested for the significance of the individual fixed effect terms with an LRT by excluding each fixed effect term at a time using the function ‘drop1’. The 95% confidence intervals for the estimates were computed using the function ‘confint’ for the GLMs or using a parametric bootstrap with 500 bootstrap samples for the GLMMs. Finally, to check which individuals had learnt and successfully transferred the concept of larger-than-large in the FTT, we checked whether each parrot chose the larger quantity more than chance using a two-sided binomial test (function ‘binom.test’ at 95% confidence level).

All figures corresponding to the statistical models were generated in R using packages ggplot2 (version 4.0.2, Wickham 2016), ggeffects (version 2.3.2, Lüdecke 2018), dplyr (version 1.2.0, Wickham et al. 2026), viridisLite (version 0.4.3, Garnier et al. 2026) and patchwork (version 1.3.2, Pedersen 2025). Figure 1 was created using Microsoft PowerPoint, with the parrot and experimenter icons generated using GPT-2.0 (OpenAI https://www.openai.com), access to which was granted by the u:ai service of the University of Vienna.

## Results

### Natural preference test (NPT)

The six parrots completed this test between 5 and 9 sessions. Initially, the parrots chose the small and large quantities at chance level (performance in the first session: Mean ± SD = 56.33% ± 18.87), indicating that they did not show any prior preference or bias at the start of our study. The parrots showed a preference for a larger quantity with increased experience across the sessions (LRT: *χ*^2^_1_ *=*12.82*, p <* 0.001; Table 1A, Figure 2a), and performed better with a larger difference between the two stimuli (*χ*^2^_1_ *=*16.45*, p <* 0.001; Figure 2b). These results confirm that the prompting of natural preference for larger quantities was successful before the basic training started.

**Figure 2:**
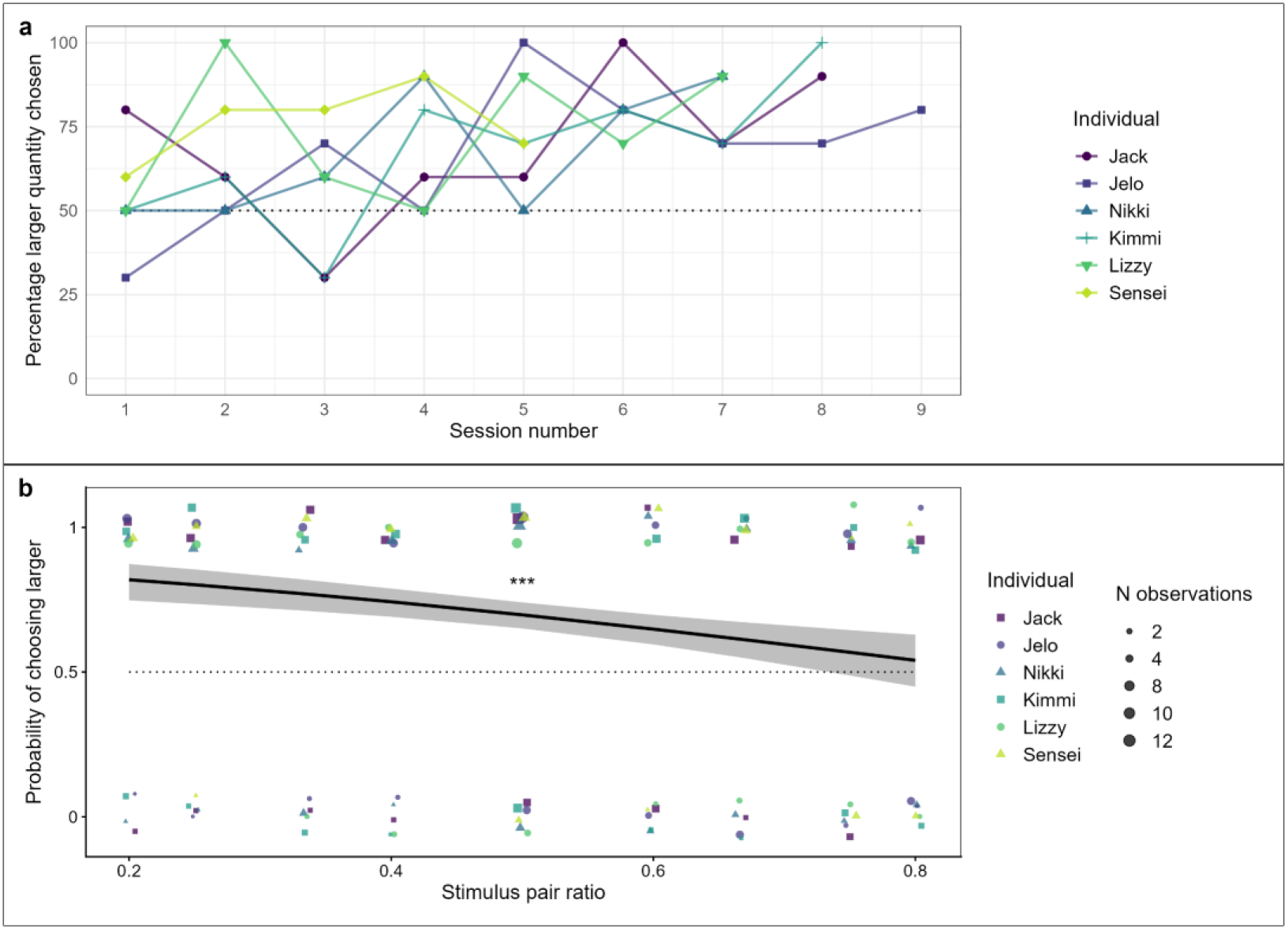
Natural preference test performance across basic training sessions for each parrot. (**a**). Probability of correct choice (choosing larger) based on the stimulus pair ratio (**b**) in the Natural preference test. The dotted lines represent the chance level. The points represent data for each individual, sizes of which are scaled by the number of observations in (**b**). The grey area represents the 95% confidence intervals around the adjusted predictions (solid black line).

**Table 1:**
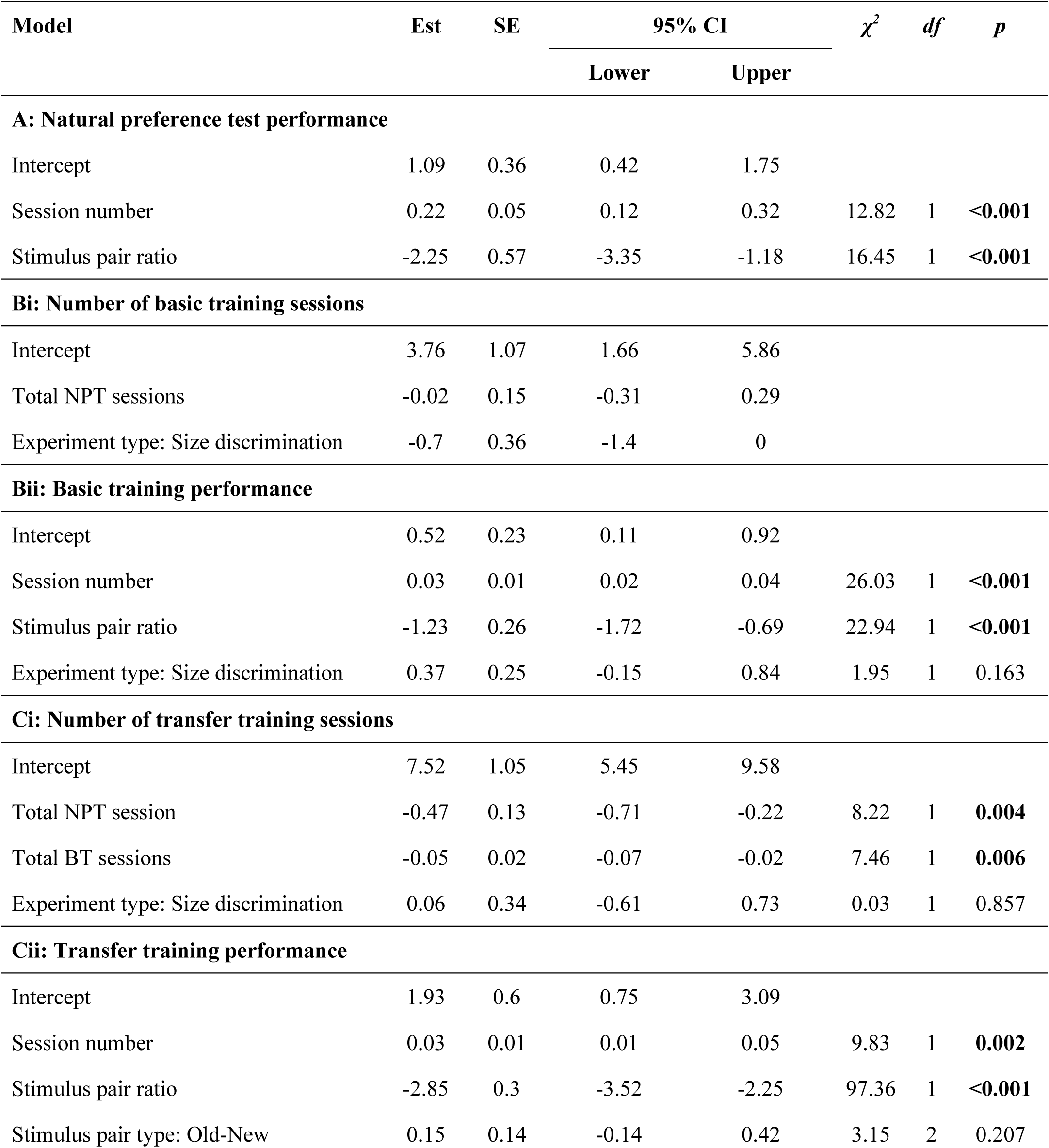
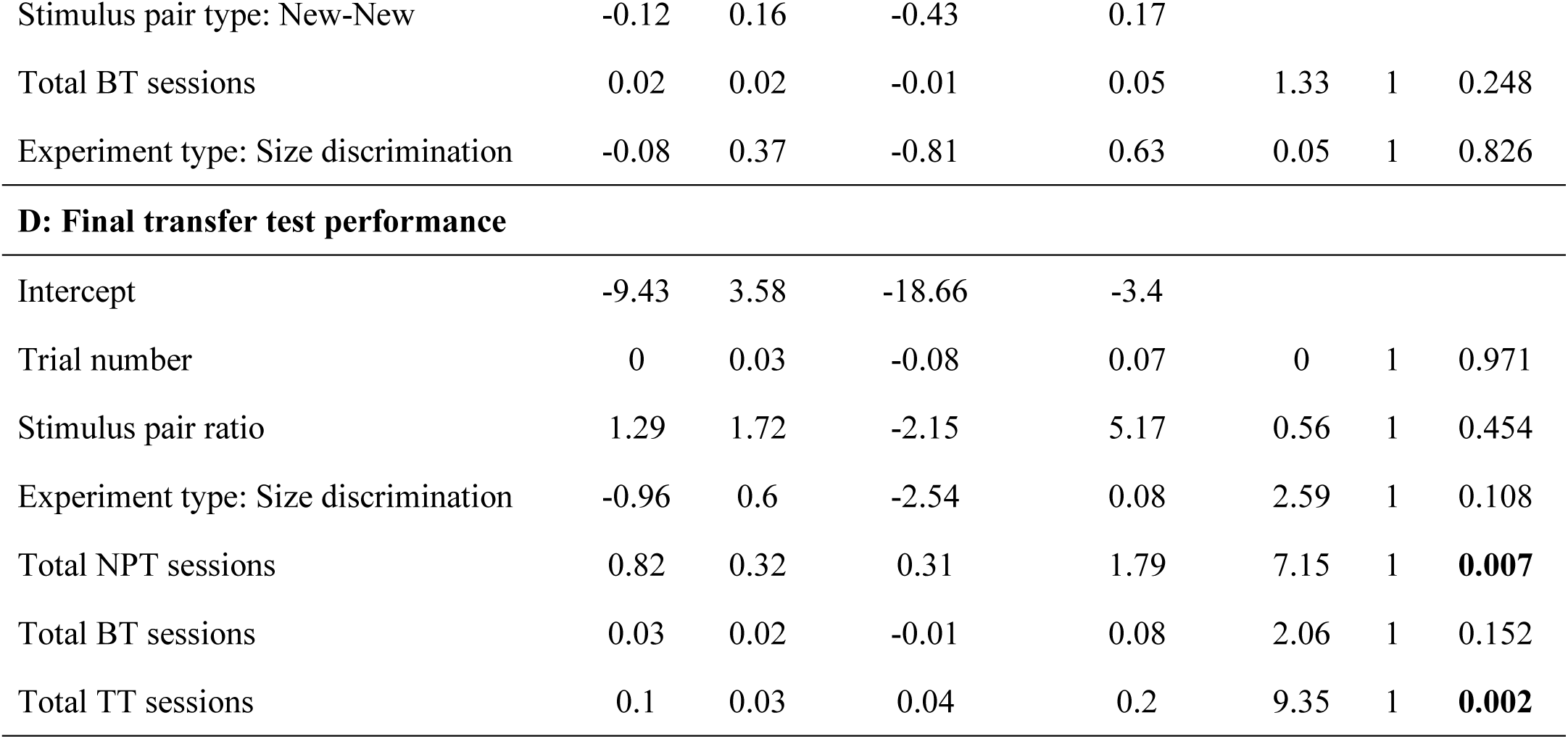
Results for models for the different phases of the study, 1A: natural preference test, 1B: basic training, 1C: transfer training, 1D: final transfer test. This table shows estimates (Est), standard errors (SE), 95% Confidence intervals (CI), ***χ***^2^, df, and p-values (bolded values indicate significant p < 0.05). Note that model tests for the significance of individual fixed predictors were conducted if the full-null model comparison was significant.

| Model | Est | SE | 95% CI | | $\chi^2$ | df | p |
| --- | --- | --- | --- | --- | --- | --- | --- |
|  |  |  | Lower | Upper |  |  |  |
| A: Natural preference test performance |  |  |  |  |  |  |  |
| Intercept | 1.09 | 0.36 | 0.42 | 1.75 |  |  |  |
| Session number | 0.22 | 0.05 | 0.12 | 0.32 | 12.82 | 1 | <0.001 |
| Stimulus pair ratio | -2.25 | 0.57 | -3.35 | -1.18 | 16.45 | 1 | <0.001 |
| Bi: Number of basic training sessions |  |  |  |  |  |  |  |
| Intercept | 3.76 | 1.07 | 1.66 | 5.86 |  |  |  |
| Total NPT sessions | -0.02 | 0.15 | -0.31 | 0.29 |  |  |  |
| Experiment type: Size discrimination | -0.7 | 0.36 | -1.4 | 0 |  |  |  |
| Bii: Basic training performance |  |  |  |  |  |  |  |
| Intercept | 0.52 | 0.23 | 0.11 | 0.92 |  |  |  |
| Session number | 0.03 | 0.01 | 0.02 | 0.04 | 26.03 | 1 | <0.001 |
| Stimulus pair ratio | -1.23 | 0.26 | -1.72 | -0.69 | 22.94 | 1 | <0.001 |
| Experiment type: Size discrimination | 0.37 | 0.25 | -0.15 | 0.84 | 1.95 | 1 | 0.163 |
| Ci: Number of transfer training sessions |  |  |  |  |  |  |  |
| Intercept | 7.52 | 1.05 | 5.45 | 9.58 |  |  |  |
| Total NPT session | -0.47 | 0.13 | -0.71 | -0.22 | 8.22 | 1 | 0.004 |
| Total BT sessions | -0.05 | 0.02 | -0.07 | -0.02 | 7.46 | 1 | 0.006 |
| Experiment type: Size discrimination | 0.06 | 0.34 | -0.61 | 0.73 | 0.03 | 1 | 0.857 |
| Cii: Transfer training performance |  |  |  |  |  |  |  |
| Intercept | 1.93 | 0.6 | 0.75 | 3.09 |  |  |  |
| Session number | 0.03 | 0.01 | 0.01 | 0.05 | 9.83 | 1 | 0.002 |
| Stimulus pair ratio | -2.85 | 0.3 | -3.52 | -2.25 | 97.36 | 1 | <0.001 |
| Stimulus pair type: Old-New | 0.15 | 0.14 | -0.14 | 0.42 | 3.15 | 2 | 0.207 |
| Stimulus pair type: New-New | -0.12 | 0.16 | -0.43 | 0.17 |  |  |  |
| Total BT sessions | 0.02 | 0.02 | -0.01 | 0.05 | 1.33 | 1 | 0.248 |
| Experiment type: Size discrimination | -0.08 | 0.37 | -0.81 | 0.63 | 0.05 | 1 | 0.826 |
| D: Final transfer test performance |  |  |  |  |  |  |  |
| Intercept | -9.43 | 3.58 | -18.66 | -3.4 |  |  |  |
| Trial number | 0 | 0.03 | -0.08 | 0.07 | 0 | 1 | 0.971 |
| Stimulus pair ratio | 1.29 | 1.72 | -2.15 | 5.17 | 0.56 | 1 | 0.454 |
| Experiment type: Size discrimination | -0.96 | 0.6 | -2.54 | 0.08 | 2.59 | 1 | 0.108 |
| Total NPT sessions | 0.82 | 0.32 | 0.31 | 1.79 | 7.15 | 1 | **0.007** |
| Total BT sessions | 0.03 | 0.02 | -0.01 | 0.08 | 2.06 | 1 | 0.152 |
| Total TT sessions | 0.1 | 0.03 | 0.04 | 0.2 | 9.35 | 1 | **0.002** |

### Basic training (BT)

All three parrots (Jack, Jelo and Nikki) who participated in the SDT task passed BT, requiring between 8 and 25 sessions. In the first session, the mean correct performance was 43.33% (± 5.77). The other three parrots (Kimmi, Lizzy, and Sensei) participated in the QDT and took 24 to 57 sessions to complete BT, with each parrot showing 60% correct performance in their first session (mean group performance = 60% correct).

The number of sessions taken in the NPT did not significantly affect the number of sessions taken to complete the BT (*χ*^2^_1_ *=* 0.01*, p =* 0.923, Table 1Bi). Whether the parrots participated in SDT or QDT also did not significantly affect performance across training sessions (*χ*^2^_1_ *=* 1.95*, p =* 0.163, Table 1Bi), indicating no difference between the two BTs. The parrots showed an overall increase in choosing the larger size or quantity across sessions in both tasks (*χ*^2^_1_ *=* 26.03*, p <* 0.001; Table 1Bii, Figure 3a). Stimulus pair ratio significantly predicted correct choices made across training sessions (*χ*^2^_1_ *=* 22.94*, p <* 0.001; Table 1Bii, Figure 3b); that is, parrots made more correct choices when a larger difference is shown in the stimulus pairs (a lower ratio).

**Figure 3:**
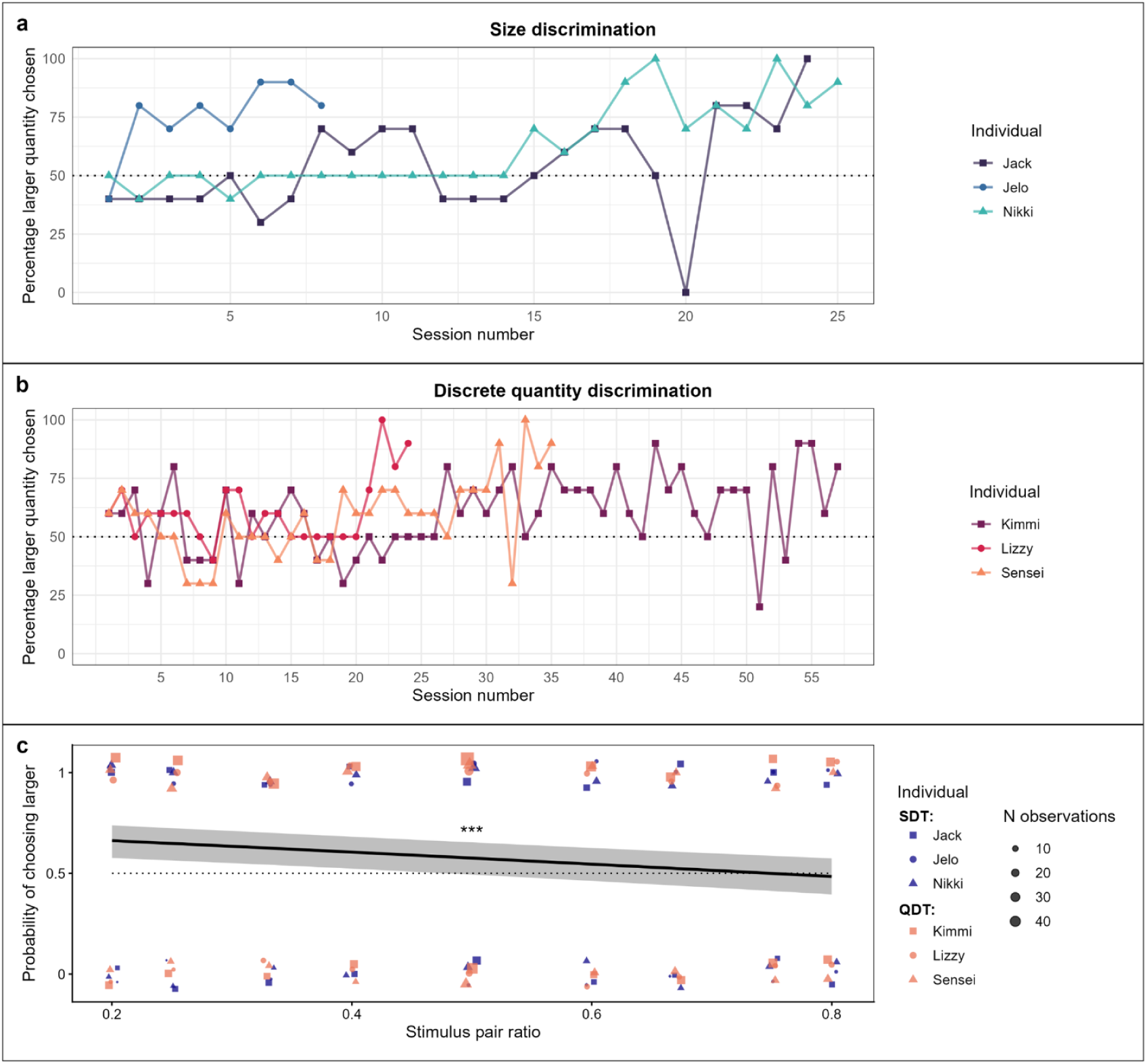
Basic training performance across sessions in (**a**) the size discrimination task (SDT) and (**b**) the quantity discrimination task (QDT) for each parrot. (**c**) The choice of the larger quantity was influenced by the stimulus pair ratio. The dotted lines represent the chance level. The points represent data for each individual, sizes of which are scaled by the number of observations in (**c**). The grey area represents the 95% confidence intervals around the adjusted predictions (solid black line).

### Transfer training (TT)

TT included three types of stimulus pairs: pairs of old-old stimulus included a change of shape in presentation, but the stimuli were within the learned range as in BT, old-new stimulus pairs included a change of shape in presentation, with one stimulus within the learned range in BT and a novel second one, new-new stimulus pairs included a change of shape in presentation and both stimuli were beyond the learned range in BT. All six parrots completed TT; the three parrots in SDT completed TT within 15 to 22 sessions, with the first TT session showing no better than chance (35-65% correct responses). For QDT, all three birds completed TT, with the first TT session showing a mean of 71.67% (± 5.77) correct responses. Specifically, Kimmi took three sessions to reach the TT training criterion, with a 79% correct response to old-new and new-new stimulus pairs on the first TT session. Lizzy took 19 TT sessions to reach the training criterion, with a 74% correct response to old-new and new-new stimulus pairs. Finally, Sensei took forty-three TT sessions to reach the training criterion, with his first TT session showing a 63% correct response to old-new and new-new stimulus pairs.

The number of TT sessions taken to reach the criterion was significantly and negatively predicted by the number of NPT sessions (*χ*^2^_1_*=* 8.22*, p =* 0.004, Table 1C, Figure 4a) and the number of BT sessions (*χ*^2^_1_ *=* 7.46*, p =* 0.006; Table 1Ci, Figure 4b). Table 1Cii (Model 5) shows the GLMM results for the performance in TT. Correct choices were predicted by an increase with more training sessions (*χ*^2^_1_*=* 9.83*, p =* 0.002, Figure 4c), and a larger difference between the stimulus pairs (i.e., lower ratio) (*χ*^2^_1_ *=* 97.36*, p* < 0.001, Figure 4d).

**Figure 4:**
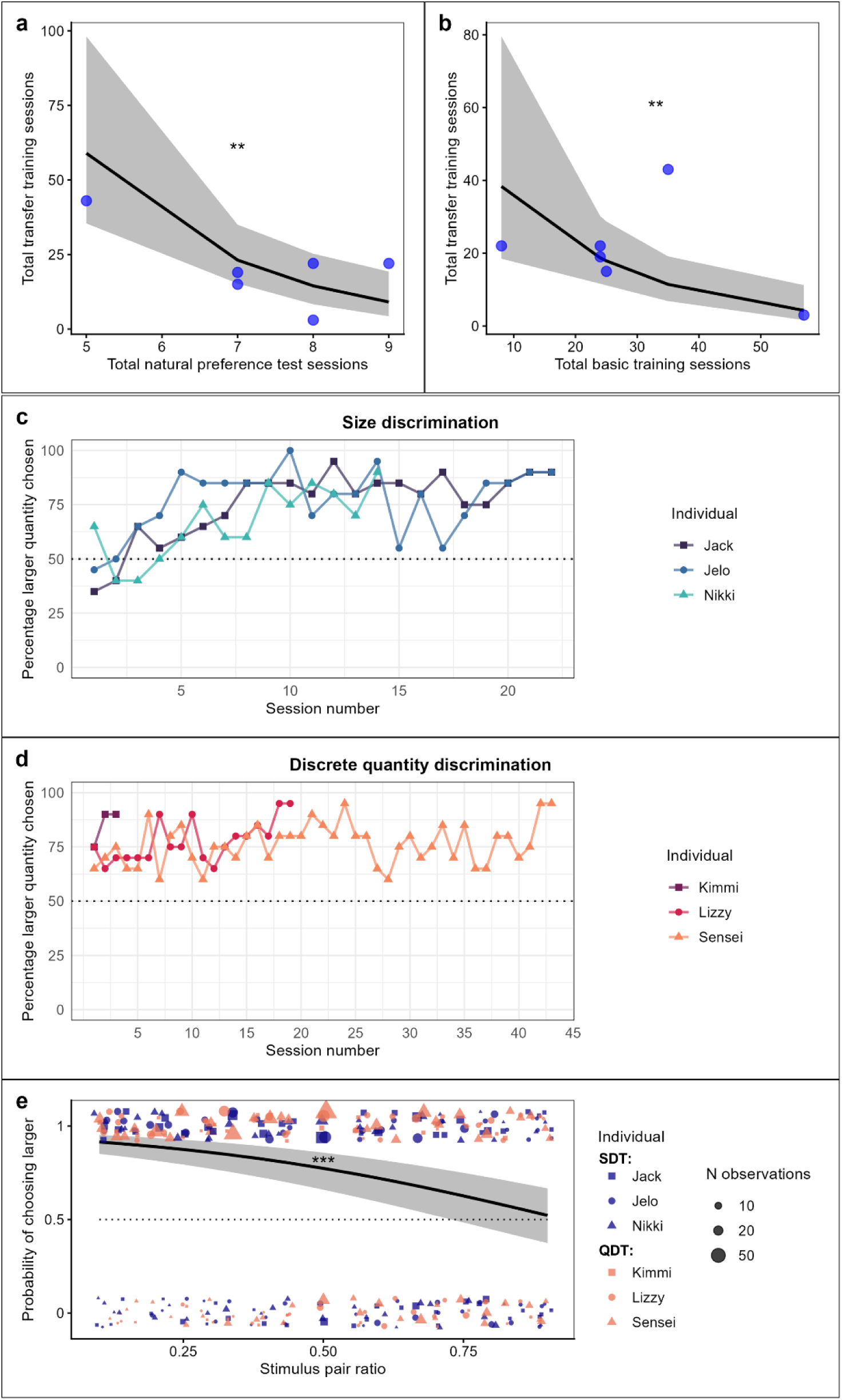
The number of sessions required to complete TT was predicted by the number of (**a**) NPT sessions and (**b**) BT sessions. TT performance across sessions for each individual in (**c**) the size discrimination task (SDT) and (**d**) the quantity discrimination task (QDT). (**e**) The choice of the larger quantity was influenced by the stimulus pair ratio. The dotted lines represent the chance level. The points represent data for each individual, sizes of which are scaled by the number of observations in (**e**). The grey area represents the 95% confidence intervals around the adjusted predictions (solid black line).

### Final transfer test (FTT)

The FTT consisted of only one session and included stimuli beyond the presentation range of BT and TT. All parrots from both SDT and QDT completed the FTT. The mean correct performance was 45% (± 21.80) in the FTT of SDT (Figure 5a), and it was 60% (± 13.23) in the QDT (Figure 5b). Importantly, one parrot, Sensei, successfully demonstrated the larger-than-large concept in the discrete quantity task, with a correct choice of 75% (binomial test: *p =* 0.041) (Figure 5b). Table 1D shows the GLMM results for FTT. Only the number of sessions taken in the NPT (LRT: *χ*^2^_1_*=*7.15*, df =* 1*, p =* 0.007, Figure 5c) and the number of sessions taken in the TT (*χ*^2^_1_*=* 9.35*, df =* 1*, p =* 0.002, Figure 5d) significantly and positively predicted correct choices of the session.

**Figure 5:**
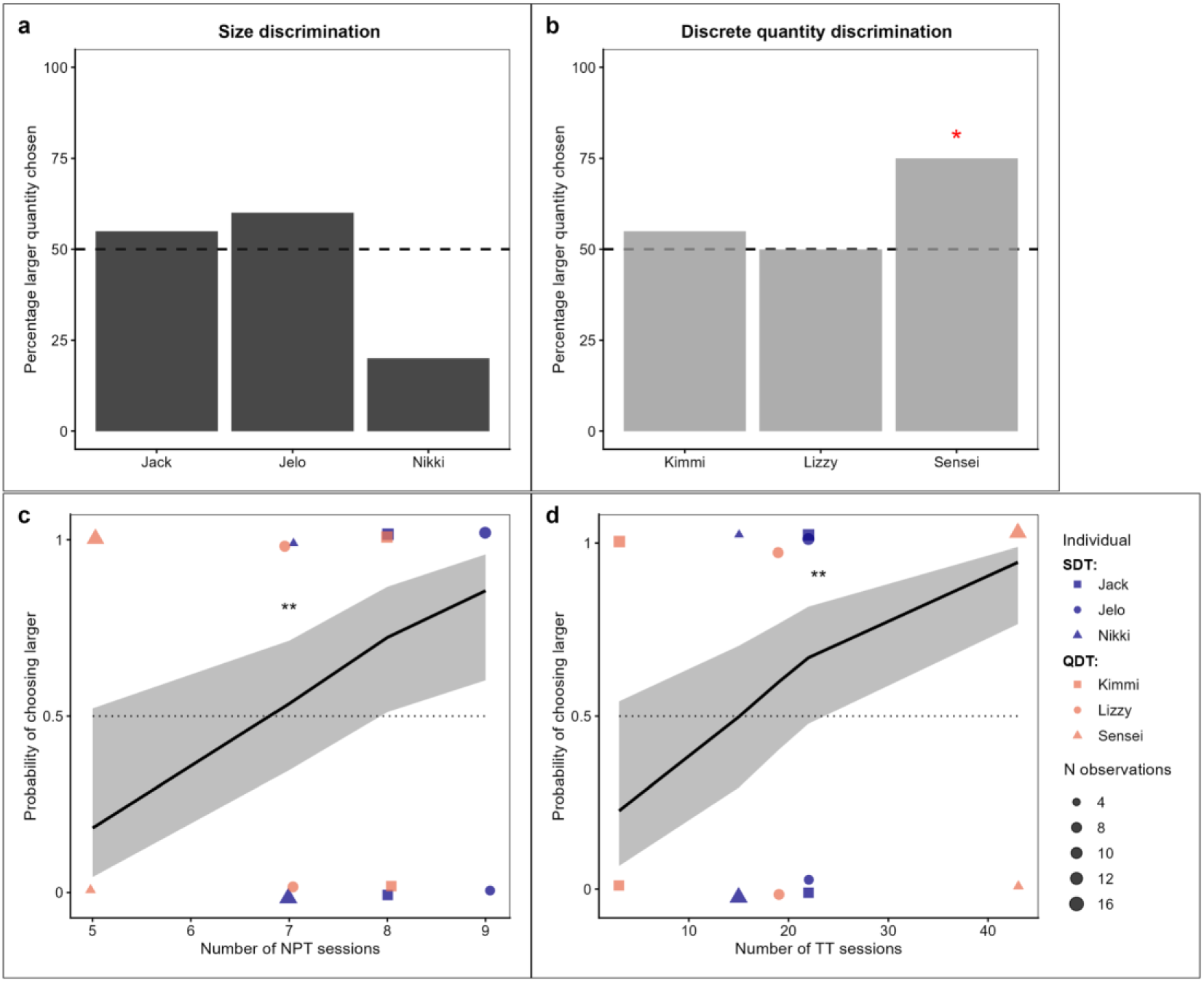
Final transfer test performance in (**a**) the size discrimination task (SDT) and (**b**) the discrete quantity discrimination task (QDT). This performance was predicted by the number of (**c**) NPT sessions and (**d**) BT sessions required. The dotted lines represent the chance level. The points represent data for each individual, sizes of which are scaled by the number of observations in (**c**) and (**d**). The grey area represents the 95% confidence intervals around the adjusted predictions (solid black line).

## Discussion

Animals frequently discriminate between stimuli based on magnitudes presented in different dimensions (e.g., size, quantity, and volume). In this study, we aimed to investigate whether African grey parrots could learn a relative rule, ‘larger-than’, and generalise it to solve new stimuli (varied in shape, size, or quantitative dimension), and if so, how they did it. Our results showed that while the size discrimination task (SDT) and quantity discrimination task (QDT) did not significantly affect parrots’ performance, none of the parrots in SDT passed the final transfer test (FTT). However, one of the six parrots, Sensei, successfully passed the FTT in the QDT, in which the stimulus pairs fell outside the quantity range used during the transfer training (TT). Sensei’s performance, therefore, provides evidence that he applied the relative larger-than rule to unseen quantities. The discussion below focuses on the possible explanations for how the parrots completed the larger-than basic training and transfer training. We also consider possible explanations for Sensei’s successful completion on the larger-than FTT.

In the natural preference test, the parrots started with a neutral preference between a smaller and a larger portion of seeds, and gradually increased their preference toward the larger portion in the test. However, this gradual shift toward the larger portion of seeds did not spontaneously facilitate the parrots choosing the larger stimulus in their first session of the BT as parrots performed at chance level. While our results are similar to previous findings that all parrots are able to discriminate between different sizes and quantities (Pepperberg and Brezinsky 1991b; Pepperberg 2006d; Aïn et al. 2009b), their performance across BT sessions may be explained by the use of a trial-and-error strategy (Thorpe 1951), as the parrots learned to choose the larger stimulus (correct choice) with repeated and accumulated experience in BT. These results highlight the importance of accumulated experience of the pairings in facilitating learning. With repeated exposures to the finite pairings in the BT, the parrots became increasingly familiar with the stimulus pairs. Such familiarity may allow the parrots to gradually form associations between the correct choice of each stimulus pair and reward (Zentall et al. 2014b).

In TT, we aimed to assess whether parrots had learned the relational rule (choose the ‘larger-than’ stimulus) in the BT, and generalised it to solve novel stimulus pairs in TT. Stimulus pairs were presented in three types in TT: old-old stimulus pairs had both stimuli presented within the range of BT, but in novel shape; old-new had one of the two stimuli presented beyond the range of BT in novel shape; and new-new stimulus pairs had both stimuli completely beyond the range of BT and in novel shape. At the beginning, we hypothesised that if the parrots had formulated the relational rule and applied it in the TT, particularly when solving the old-new and new-new stimulus pairs, then performance was expected to be beyond chance level. However, as a group, the parrots also did not perform better than chance, and the type of stimulus pairs was not a significant predictor of correct choices.

Therefore, it seems we have limited evidence to support that the parrots had formed a relational rule during BT and systematically applied it to solve the different types of stimulus pairs in TT. At the individual level, given that two parrots in the QDT achieved 74% or above correct performance on their first session, results may suggest there is a task difference.

However, task type is not a significant predictor for choice, and thus, the results instead suggest that there is individual variation in acquiring the relational rule. These results of TT are similar to the results in BT in that they suggest that most parrots’ performance in the TT appeared to be explained by trial-and-error as well as associative learning through accumulated experience (Katz et al. 2007).

Despite this, parrots appeared to be also using another strategy, that is, discriminating stimuli based on magnitude differences in the stimulus pair. Our results showed that the parrots were more likely to choose the larger stimulus when there was a larger difference between the stimuli. Increased correct responses in relation to the proportional magnitude difference between stimuli can be explained by Weber’s law. This law states that discrimination between stimuli becomes easier as the difference between stimuli increases, which has been shown in African grey parrots (Aïn et al. 2009b) and other animal species (Schaffer et al. 2025; Worsley et al. 2025b) when solving this kind of discrimination task. Because the ratio between the two stimuli is not only a significant predictor of performance in TT, but also of performance in NPT and BT, these findings suggest that the parrots were sensitive to the relative magnitude differences between the stimuli, in addition to associative learning.

At the session/training level, the parrots showed better performance with increased session number, and the parrots that had more BT, alongside their gradually increasing preference in NPT, needed fewer sessions to reach the learning criteria of TT. Considering performance at the trial level, as discussed above, one of the possible explanations for the increase in performance with more sessions is that more sessions may allow the parrots to increase familiarity with the stimulus pairs as well as the reward contingencies. Such familiarity may facilitate the formation of the relational rule (i.e., choose the larger stimulus) during the BT, allowing the parrots to apply this rule more efficiently when solving the novel stimulus pairs (i.e., the old-new and new-new stimulus pairs) in TT (Zentall et al. 2014b). Greater experience in transfer training, but not in the BT, also led to better performance in the FTT. This may suggest that experience with different types of stimulus pairs during TT was related to subsequent FTT performances through the old-new and new-new stimulus pairs. As a group, most parrots failed the FTT except one, Sensei, who chose the larger stimulus above chance, thus providing preliminary evidence consistent with the ability to generalise the relational rule to novel stimulus pairs. Sensei had at least twice as many TT sessions as the other two parrots in the QDT. Because Sensei had the fewest NPTs to reach the criterion (i.e., quickly demonstrated a preference for the larger portion of seeds in the NPT), Sensei’s quicker acquisition of the larger preference, in addition to the extensive training, may have facilitated subsequent learning.

As mentioned before, the concept of larger-than exists in different magnitudes beyond size and quantity tested here, and thus, ideally the parrots could be tested across different magnitudes (e.g., SDT→QDT or vice versa) to demonstrate that they could acquire the full abstract concept (Flemming and Beran 2012). However, we were only able to assess one of the two dimensions for each parrot due to limited time and resources available during COVID-19. Therefore, whether the parrots are better at demonstrating the understanding of one dimension than another remains to be investigated (Pepperberg 2020). While none of the parrots in the SDT passed the final transfer test, a previous study with a single African grey parrot, Alex, showed the ability to generalise the relative-size concept to new stimuli (Pepperberg and Brezinsky 1991b), and other studies have shown African grey parrots’ understanding of ordinal relationships in quantity (Pepperberg 2006d; Aïn et al. 2009b). Therefore, these results and previous findings suggest that the extent to which one grasps the concept varies individually in grey parrots.

Understanding the concept of relativity is one step above simple associations, as individuals extract a relational rule from what is previously known, learned, or experienced. Such a concept has been related to ecological implications (Nieder 2020b), for example, ‘larger-than’ may be useful for maximising foraging when comparing available options. Our results here indicated that most parrots do not necessarily need to demonstrate a true understanding of ‘larger-than’. Despite this, having such an representation of environments may allow for faster or more flexible adjustments when environmental conditions change. For example, such representation could be advantageous for continued exploration, moving between locations, and selecting resources beyond those previously experienced. Therefore, understanding such representational concepts nevertheless contributes to the understanding of how animals navigate and make choices in their environment and whether forming these concepts would require more advanced cognitive processes.

## Author Contributions

P.K.Y.C.: Conceptualization, Data curation, Formal analysis, Funding acquisition, Investigation, Methodology, Project administration, Supervision, Writing - original draft, Writing - review and editing A.B.: Conceptualization, Data curation, Formal analysis, Investigation, Methodology, Visualization, Writing - original draft, Writing - review and editing J.R.D.: Validation, Writing - review and editing T.R.: Conceptualization, Methodology, Writing - review and editing A.M.P.vB.: Resources, Writing - review and editing

## Funding

This research was funded by Animal Mind Project to P.K.Y.C.

## Institutional Review Board Statement

Following the German Animal Welfare Act of 25 May 1998, Section 5, Article 7, and the Spanish Animal Welfare Act 32/2007 of 7 November 2007, Preliminary Title, Article 3, this study was classified as a non-animal experiment, and thus it did not require approval from a relevant body. The care and use of animals were strictly adhered to the ethical guidelines of the Association for the Study of Animal Behaviour and Animal Behaviour Society.

## Data Availability Statement

Data are uploaded to OSF for review and are publicly available for the public (https://doi.org/10.17605/OSF.IO/QVNW5).

## Acknowledgements

We thank the president of the Loro Parque, Wolfgang Kiessling, and his staff for their generous support and for providing us with access to the birds and the research facilities. We thank the Loro Parque Fundación and its president, Christoph Kiessling, for their collaboration and the staff of the Loro Parque Fundación for their support. We thank A Krasheninnikova for general advice and feedback on lab logistics.

## Conflicts of Interest

All authors declare that there is no conflict of interest.

## Notes

https://doi.org/10.17605/OSF.IO/QVNW5

### Competing Interest Statement

The authors have declared no competing interest.

